# Not two sides of the same coin: Divergent effects of motor state on corticospinal and cortical responses to TMS of motor cortex

**DOI:** 10.1101/2025.09.07.674734

**Authors:** Mikkel Malling Beck, Lasse Christiansen, Marieke Heyl, Angela Mastropasqua, Sybren Van Hoornweder, Axel Thielscher, Leo Tomasevic, Hartwig Roman Siebner

## Abstract

**Background:** Transcranial magnetic stimulation (TMS) of the motor cortex can elicit motor evoked potentials (MEPs) in target muscles, reflecting corticospinal excitability. MEP amplitudes increase with TMS intensity and can be facilitated by tonic muscle pre-activation. Since conventional transcranial evoked potentials (TEPs) also grow with increasing TMS intensity, cortical and corticospinal responses are often considered two facets of the same process. If this were true, changes in physiological motor state should modulate TEPs and MEPs in a similar manner.

**Methods:** To compare the state-dependency of cortical and corticospinal responses to single-pulse TMS, we simultaneously recorded TEPs and MEPs in 16 healthy young adults during relaxation and isometric contraction of the right first dorsal interosseous (FDI) muscle. For each condition, 100 biphasic TMS pulses were delivered to the left primary motor hand area at five different intensities centered around the resting motor threshold.

**Results:** TEP and MEP amplitudes increased with stimulation intensity. As predicted, tonic muscle contraction consistently facilitated MEP. On the contrary, muscle contraction attenuated two key peaks of the TEP (N15 and N100). The state-dependent effects of corticospinal and cortical responses were not correlated.

**Discussion:** Both TEPs and MEPs are reliably modulated by motor state, yet they differ in direction and their magnitudes do not scale with each other. These findings challenge the assumption that cortical and corticospinal responses are two aspects of the same process. MEP facilitation during contraction likely reflects increased spinal excitability, whereas TEP attenuation may reflect reduced responsiveness of cortico-cortical or cortico-subcortical networks.

## Introduction

Transcranial magnetic stimulation (TMS) uses electromagnetic induction to non-invasively excite neural tissue in the human cortex. The direct neural excitation generated by a single TMS pulse in the cortical target site propagates to connected cortical, subcortical and spinal structures and back causing both local, remote, and reverberant activity [1]. TMS applied over the primary motor hand region (M1-HAND) at sufficient intensities creates a volley of action potentials that travel down the corticospinal tract and activate spinal motoneurons to produce a motor evoked potential (MEP) in the targeted hand muscle measured via electromyography (EMG). The possibility to combine TMS with electroencephalography (EEG) [2,3] enables a direct characterization of cortical responses to stimulation [3]. The TMS-evoked potential (TEP) consists of a series of positive and negative peaks or deflections in the averaged EEG time-series [4–6] that result from the activation of multiple neural sources with distinct spatiotemporal profiles.

While MEPs and TEPs are both measuring physiological responses to TMS, they may represent different neurophysiological phenomena. The MEP is a peripheral motor readout reflecting the excitability and signal propagation along the entire corticospinal pathway. This includes the targeted cortical circuitry in the precentral cortex, the corticospinal projections, the spinal motoneuron pool and possible interneuron relays and the peripheral motor axon [1,7]. Hence, alterations in the state of the corticospinal pathway can alter MEP amplitudes without this necessarily implying an excitability change within the initially stimulated cortical volume. On the other hand, the mechanisms generating the characteristic TEP peaks – N15, P30, N45, P60 and N100– remain unknown. Given their latency, occurring more than 10ms after the TMS pulse, these TEP peaks likely do not reflect the initial activation of principal cells or inhibitory interneurons by the TMS-induced electric field in the motor cortex that underlies MEP generation. Instead, they may reflect reverberations of excitability changes within activated cortico-subcortical circuits. This implies that MEPs and TEPs may not respond in a similar manner to “extrinsic” TMS variables like stimulus intensity, or “intrinsic” brain variables such as changes in brain states.

A characteristic feature of MEPs is that their amplitudes increase with increasing TMS intensities and with voluntary pre-activation of the target muscle [8,9]. TEP peak amplitudes have also been shown to increase with stimulus intensity [6,10,11]. The comparable dose-response relationship is compatible with the idea that corticospinal and cortical responses are just different aspects of the same underlying phenomenon. If this was the case, TEPs and MEPs should also share a similar sensitivity to a change in physiological state. A prominent state-dependent feature is a consistent and prominent facilitation of MEP amplitudes during voluntary pre-activation of the target muscle relative to the MEP evoked during muscle relaxation and can be largely attributed to spinal mechanisms [12].

This prominent example of state dependency motivated this investigation into whether voluntary motor activity differentially modulates cortical and corticospinal responses to TMS in healthy volunteers. We simultaneously recorded MEPs and TEPs across a range of stimulus intensities during two physiological states, namely while participants were at rest (i.e., relaxed target muscle) and performed a tonic contraction (i.e., steady muscle engagement). The simple state manipulation revealed distinct modulations of specific cortical and corticospinal responses, indicating that TEPs and MEPs reflect separate, not equivalent, aspects of TMS-induced neurophysiological activity.

## Methods

### Participants

Twenty-eight healthy volunteers (10 males; age range: 23-32 years; mean age = 26.3 ± 1.9 years) with no history of neurological or psychiatric disorders participated in the experiment after giving informed consent, receiving written information, and being screened for magnetic resonance imaging (MRI) and TMS contraindications. All experimental procedures were carried out in accordance with the Helsinki declaration and were approved by the regional ethical committee (Capital Region of Denmark; Protocol number H-15008824).

### Transcranial magnetic stimulation (TMS)

Single-pulse TMS was delivered every 2 seconds (jitter ± 0.2 seconds) to the hand representation of the left primary motor region (M1-HAND) using biphasic pulse waveforms (MagVenture X100 with MagOption, MagVenture A/S, Farum, Denmark) and a 70-mm figure-of-eight coil (MC-B70 coil, MagVenture A/S, Farum, Denmark). Neuronavigation (TMS Navigator, Localite GmhB, Bonn, Germany) was used to monitor the coil position throughout the experiment using individual T1w structural images (3T MRI, Siemens PRISMA, Erlangen, Germany). The precentral stimulation site was determined using an individualized mapping procedure (for similar approach, see [13]). First, the precentral motor “hand knob” was anatomically localized [14]. Then, a mini-mapping procedure around this spot was performed to localize the MEP hotspot, i.e., the spot with most consistent and largest motor evoked potentials (MEPs) in the first dorsal interosseus (FDI) muscle. Next, the resting motor threshold (RMT) was estimated defined as the intensity eliciting MEPs >50 µV in 5/10 stimuli [15]. After having identified the individual MEP hotspot and the RMT, EEG data were inspected using the rt-TEP tool [16] to evaluate whether scalp muscle artefacts were triggered. Scalp muscle artefacts following TMS of M1-HAND are characterized as biphasic responses with topographic extremes in EEG electrodes close to the temporal muscle [13,17]. If scalp muscle responses were observed, the coil was moved or tilted away from the activated temporal muscle. This successfully eliminated scalp muscle artefacts in 16 of the 28 participants. In the remaining 12 participants,minor coil adjustments failed to avoid muscle artifacts, and the experiment was discontinued.

Subsequently, TMS was delivered at 5 intensities and during 2 motor states. Per intensity and motor state, 100 stimulations were delivered, resulting in 1,000 pulses per participant. Intensities were defined in relation to the RMT by changing the stimulator output in steps of 4% maximum stimulator output (%MSO): -8% MSO; -4% MSO; RMT, +4% MSO; +8% MSO. TMS was delivered while participants were at rest or while they performed a sustained isometric voluntary contraction of the index finger of the right hand at 10% of maximum voluntary contraction (MVC). Feedback was provided in the form of a rectified and smooth EMG trace from the FDI (Figure 1).

**Figure 1.**
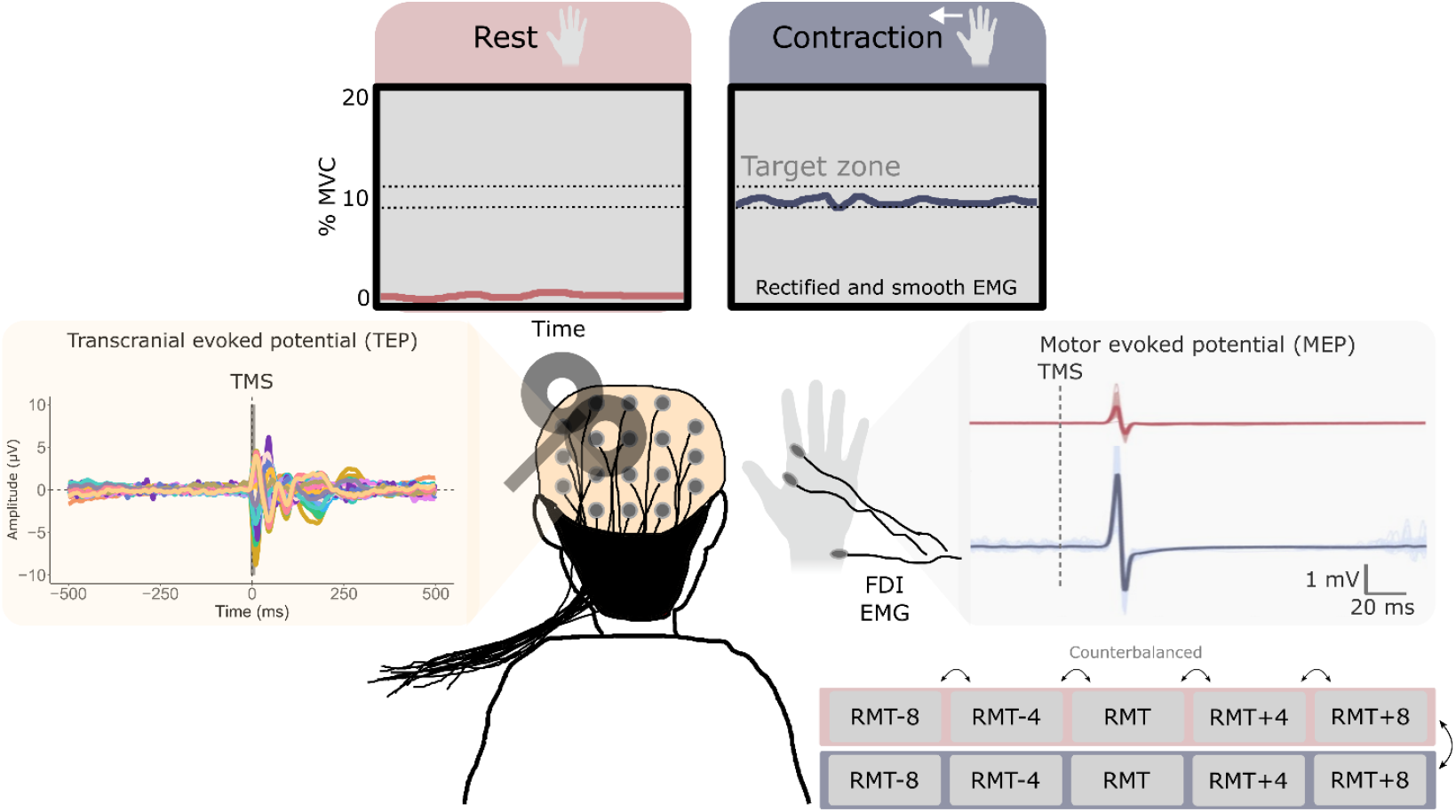
Experimental setup and measurements. Single biphasic TMS pulses were delivered at the optimized hot spot of the M1-HAND in the absence of TMS induced cranial muscle artifacts. TMS was given at five intensities while participants were at rest or performed a voluntary isometric contraction of the index finger. The cortical (TEP) and corticospinal responses (MEP) to TMS were recorded. Abbreviations: FDI = First dorsal interosseous; RMT = Resting motor threshold; MVC = Maximal voluntary contraction. RMT = Resting motor threshold.

During recording blocks, participants were asked to keep their eyes open, avoid blinking, and fixate their gaze at a fixation spot on the screen approximately 1 m in front of them. Participants were equipped with modified earplugs playing a masking sound consisting of pink noise with added TMS clicks. The masking sound was generated using TAAC software [18] and the sound pressure was adjusted so that participants reported not being able to hear the sound of the discharging TMS coil or that it was greatly reduced while the coil was floating over the head of the participants. Additionally, a thin layer of foam (approx. 1-2 mm) was used between the coil and the EEG cap to reduce bone conduction of sound and vibration from the discharging coil [19]. After each recording block, participants were asked to rate their perceived TMS sensations on a numerical rating scale ranging from 0-10. Specifically, participants rated the audibility of the TMS “click” sound; the sense of focality or “spread” of the TMS pulse on the head; the vibrations caused by the discharge of the coil; and the pressure of the coil on the head.

### Electroencephalographic (EEG) and electromyographic (EMG) recordings

EEG activity was recorded at a sampling rate of 5 kHz from 61 passive Ag/AgCl C-slit electrodes placed in an equidistant EEG cap (M10 cap layout, BrainCap TMS, Brain Products GmbH, Germany) using a TMS-compatible EEG amplifier (BrainAmp DC, Brain Products GmbH, Germany) and BrainVision Recorder software. The reference and ground electrodes were placed on the right and left side of the participants’ forehead, respectively. Electrodes were prepared using electroconductive and abrasive gel to lower the electrode impedance (<5 kΩ). Electrode impedance levels were checked regularly throughout the experiment. EMG was acquired from the right first dorsal interosseus (FDI) using a belly-tendon montage with the ground electrode positioned on the right processus styloideus ulnae. EMG signals were amplified (x500), bandpass filtered (10-2000 Hz), and sampled at 2000 Hz (Digitimer D360, Digitimer Ltd., Hertfordshire, UK) using Signal software (v4.17, Cambridge Electronic Design, Cambridge, UK).

### EEG preprocessing

EEG preprocessing was carried out in EEGLAB (v. 2021) running in MATLAB (v. R2020a) using functions from the TMS-EEG signal analyser (TESA)[20] toolbox. First, the TMS pulse artifact was removed by replacing the signal from -2 to 8 ms around the stimulation with interpolated data points using cubic interpolation. Subsequently, EEG data was band-pass filtered from 1-48 Hz using a 2^nd^ order Butterworth filter and epoched from -500 to 500 ms around stimulation. Following this, data were visually inspected and electrodes displaying excessive noise were removed (median of 2 channels excluded per participant). Missing channels were interpolated using spherical interpolation. Trials containing eye blinks or excessive noise were removed based on visual inspection of signal amplitudes and topographies (median of 11 epochs excluded per condition). Baseline correction was applied from -205 to -5 ms to the stimulation.

### EEG data analysis

Two measures were computed to quantify TMS-evoked cortical activity. Global mean field power (GMFP) [21] was computed to extract a single reference-free cortical response measure. For the analysis of TEP peak amplitudes, we focused on data from a single electrode close to site of stimulation characterized as one of the most responsive when scrutinizing the TMS-evoked EEG topographies (Fig 2B). Peaks were estimated as the largest absolute value within pre-specified time windows for the prototypical TEP peaks following stimulation of M1 [4,6,22], N15 (10-20 ms), P30 (15-45 ms), N45 (30-60 ms), P60 (45-75 ms) and N100 (75-125 ms). EEG spectral power in the time-period leading up to stimulation (-500 to -10 ms) was also computed using Welch’s method during both rest and contraction. This was done to obtain an electrophysiological validation of our state-modulation, as desynchronization of alpha and beta power is a hallmark EEG feature accompanied by motor activity [23], and to evaluate whether individual modulations of alpha and beta power during voluntary sustained contractions were associated with individual differences in TEP and MEP amplitudes. We computed the area under the power spectrum curve for the alpha-band (8-12 Hz) and beta-band (15-35 Hz) to estimate modulation of band-limited power during movement execution.

**Figure 2.**
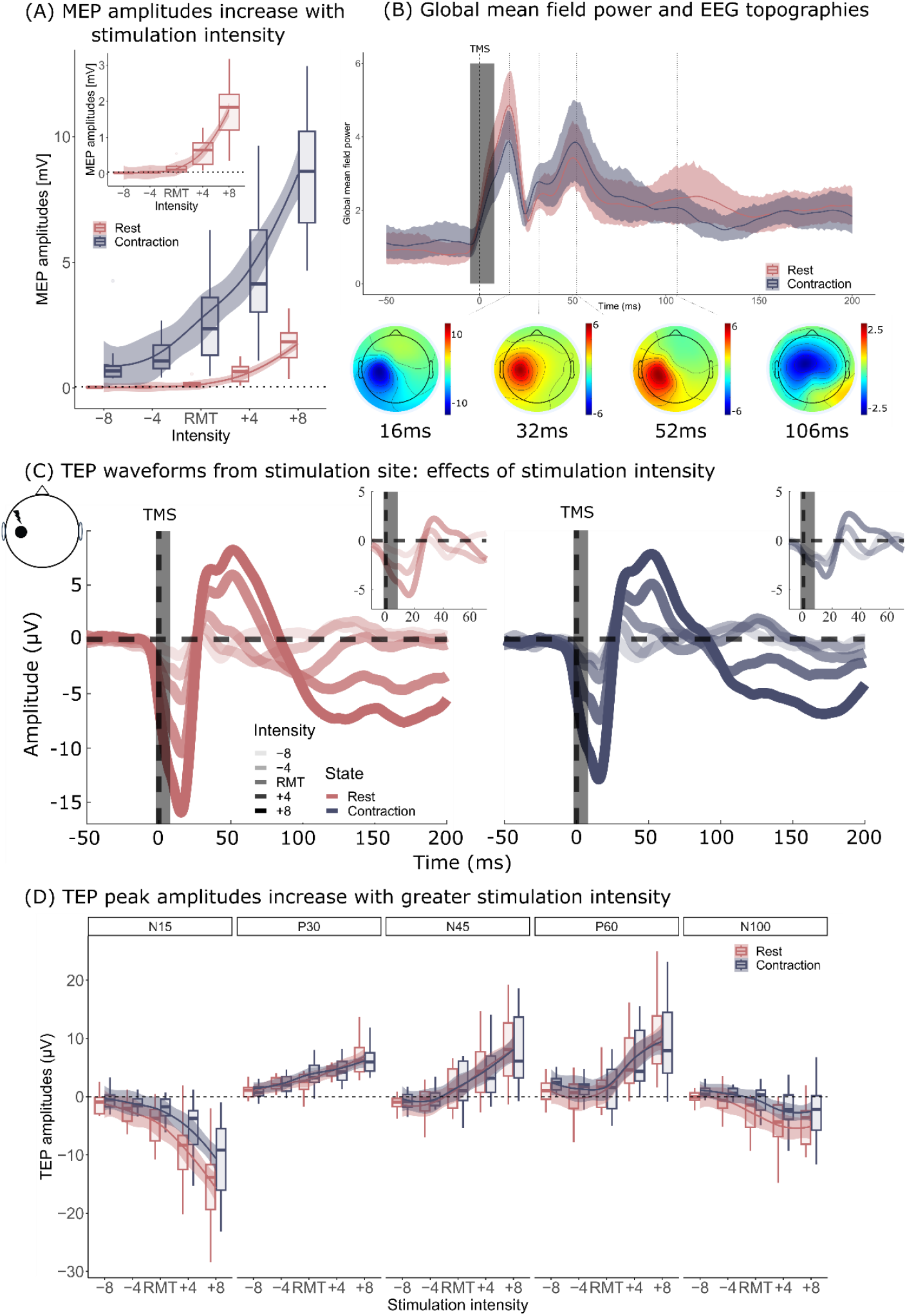
Motor evoked potentials (MEPs) and transcranial evoked potentials (TEPs) increase with stimulation intensity. (A) Effects of stimulation intensity on MEP amplitudes. (B) Average global mean field power (GMFP) and EEG topographies during rest and contraction. (C) Averaged local TEP waveforms across stimulation intensities during rest and contraction. (D) Averaged amplitudes for prototypical TEP peaks across stimulation intensities during rest and contraction.

### Statistical analysis

Two separate linear mixed effect models were used to model the effects of stimulation conditions for the extracted MEP and TEP amplitudes, respectively.

For TEPs, a mixed model was fitted to the extracted amplitudes with an interaction between three independent factors: peak (5 levels: N15, P30, N45, P60, N100); intensity (5 levels: -8%, -4%, RMT, +4%, +8%) and motor state (2 levels: rest vs. contraction). Individuals (ID) were added to the model as random intercepts:

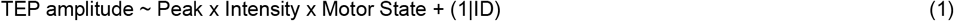

Where *x* represents an interaction and (1|ID) represents the random intercepts for each participant.

A similar model was fitted for the MEP amplitudes, but without the interaction with Peak. Furthermore, MEPs were log-transformed to adhere to assumptions of normality of residuals.

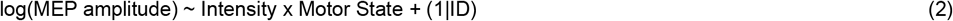

Mixed effect models were fitted in R using the *lme4-package* [24]. F-statistics and p-values for main effects and interactions were estimated using the *lmerTest-package* [25] and post-hoc comparisons were performed using the *emmeans-package* [26]. In addition, Pearson correlation analyses were performed to investigate potential associations between state-dependent modulations of TEPs, MEPs, and alpha/beta power. P-values were adjusted for multiple comparisons using the Bonferroni correction unless otherwise stated. For all analyses, alpha was set to 0.05.

## Results

### Impact of stimulation intensity of MEP and TEP peak amplitudes

TMS over M1-HAND evoked MEPs at threshold and suprathreshold intensities at rest, while TMS also evoked MEPs at subthreshold intensities during voluntary contraction. MEP amplitudes increased with stimulation intensity during both rest and contraction (Figure 2A). This was reflected by a significant main effect of stimulation intensity in the fitted mixed effect model (F = 105.1; P < 0.001). GMFPs revealed peaks at various time points during both rest and contraction (Figure 2B). The topographical maxima of activation were located close to the site of stimulation in the left sensorimotor cortex. When comparing the TEP waveforms extracted from electrodes close to this area, a general increase in peak amplitudes were observed with stimulation (Figure 2C). This was confirmed statistically by comparing the extracted peak amplitudes revealing an interaction between stimulation intensity and peak (F = 31.4; P < 0.001) (Figure 2D). Although all peak amplitudes increased with intensity, the interaction indicated that intensity had a differential effect across different TEP peaks (Figure 2D).

### Impact of voluntary motor activity on MEP and TEP peak amplitudes

As expected, MEP peak-to-peak amplitudes were consistently greater during sustained voluntary contraction compared to rest (Figure 3A). The fitted mixed effect model revealed a significant main effect of state for MEP amplitudes (F = 401.8; P < 0.001). In contrast, GMFP of the TMS evoked EEG response was not enhanced by isometric tonic contraction of the target muscle. Two components of the TEP were actually smaller during contraction than at rest (Figure 2B), namely the negative peaks around 15 ms (N15) and 100 ms (N100) after the pulse. The linear mixed model showed a significant interaction between the motor states and the extracted TEP peak amplitudes (F = 4.04; P = 0.001). Post-hoc comparisons confirmed that the state-dependent differences in TEP amplitudes were driven by an attenuation (i.e., less negativity) of the N15 peak (β_rest vs. contraction_= -2.40 ± 0.61 µV; t = 4.62; P < 0.001) and N100 peak (β_rest vs. contraction_= -1.58 ± 0.61 µV; t = 3.28; P = 0.01). No significant differences were observed for other peaks of interest (p-values range: 0.54-0.89). The state-dependent difference in N15 and N100 amplitudes did not depend on stimulation intensity as demonstrated from the lack of a three-way interaction between intensity, state, and peak (F = 0.67; P = 0.83) (Figure 3A). Together, these results show that MEP amplitudes increased during voluntary contraction, whereas amplitudes of the N15 and N100 TEP peaks decreased during voluntary contraction compared to muscle relaxation.

**Figure 3.**
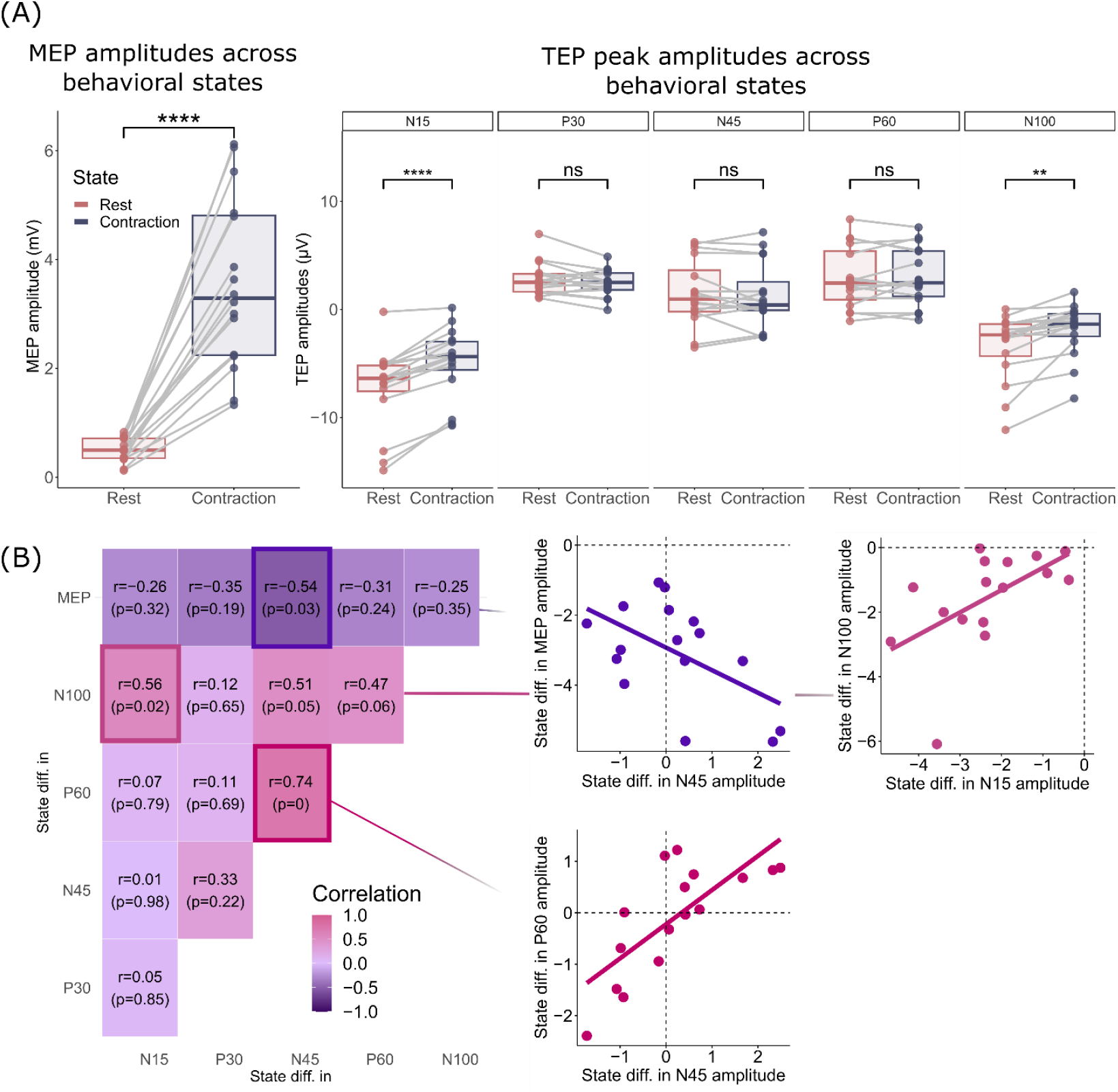
Effects of motor state on motor evoked potentials (MEPs) and transcranial evoked potentials (TEPs). (A) Amplitudes of MEPs (left) and prototypical local TEP peaks (right) during rest (red) and contraction (blue) averaged across stimulation intensities. MEP amplitudes were consistently greater during contraction compared to rest, whereas amplitudes of the N15 and N100 peaks were consistently smaller during contraction compared to rest. (B) Correlations between individual differences in TEP peak amplitudes between rest and contraction. Significant correlations were observed between the differences in the N15 and N100 amplitudes, between the N45 and P60 amplitudes and between the N45 and MEP amplitudes. P-values are corrected for the false-discovery rate.

To investigate potential associations between individual state-dependent changes in TEP peaks and MEP amplitudes across stimulation intensities a Pearson correlation matrix was computed. Significant correlations were observed between the individual differences of the state difference in the N15 and N100 amplitude (*r* = 0.56; P = 0.02), the N45 and P60 amplitude (*r* = 0.74; P = 0.002), and the N45 and MEP amplitude (*r =* -0.54, P = 0.03). These correlations are illustrated in Figure 3B.

Pre-stimulus EEG power in the alpha (8-13 Hz) and beta (15-35 Hz) frequency bands were also modulated by motor state (Figure 4A). A significant desynchronization of the alpha (t(31) = 4.47; P<0.001) and beta bands (t(31) =2.86; P=0.007) occurred during voluntary contraction compared to rest. This desynchronization was most apparent in the electrodes covering the bilateral sensorimotor cortices (Figure 4A). This indicates that our behavioral manipulation effectively altered sensorimotor brain activity. No significant correlations were observed between pre-stimulus power changes in the alpha or beta EEG bands, extracted from the electrode closest to the stimulated target region, and changes in TEP peak amplitudes for the N15 and N100 peak across stimulation intensities (Figure 4B). Correlation coefficients and p-values ranged from 0.027-0.35 and 0.18-0.92 (uncorrected), respectively.

**Figure 4.**
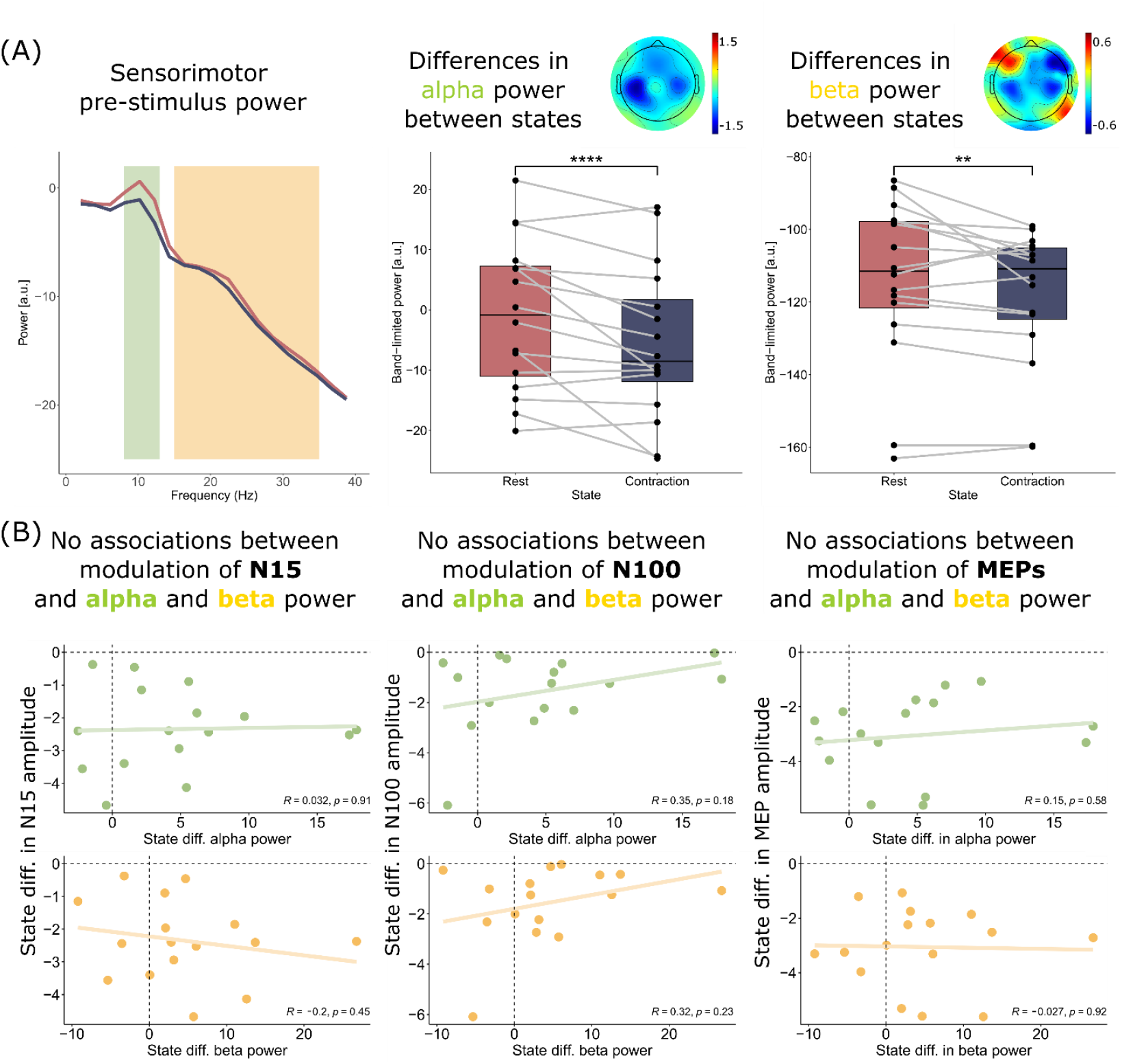
Effects of motor state on pre-stimulus alpha and beta power and associations to TEPs and MEPs. (A) Power spectrum estimated from electrode overlying the stimulated sensorimotor cortex during the pre-stimulus period. Differences in alpha (middle) and beta (right) power during rest vs. contraction and corresponding topographical plots showing differences in band-limited power across states across the scalp. (B) Individual differences in alpha and beta power were not associated with individual state-differences in N15, N100 or MEP amplitudes.

## Discussion

This study directly compared the effects of tonic muscle contraction on two distinct responses to single-pulse TMS of the M1-HAND region. Across multiple stimulation intensities, we simultaneously recorded corticospinal motor evoked potentials (MEPs), evoked by transsynaptic activation of the corticomotor pathway, and cortical transcranial evoked potentials (TEPs), reflecting the brain’s direct response to TMS. Both responses scaled with stimulation intensity but were differentially modulated by voluntary muscle activation. MEP amplitudes were consistently enhanced during tonic contraction, whereas the early (N15) and late (N100) peaks of the TEP were reliably attenuated. These findings demonstrate that TEPs and MEPs capture distinct facets of state-dependent TMS-induced activation of the motor system.

### Effect of isometric tonic contraction on the transcranially evoked response

Alterations in the brain’s functional state can markedly influence its responsiveness to TMS. A prominent example is the transition from wakefulness to slow-wave sleep, causing pronounced changes in the spatiotemporal dynamics of the TEP [27–29]. In contrast, the manipulation of motor state investigated in the present study elicited only modest, yet highly consistent, changes in the TEP magnitudes. Specifically, the transitioning from rest to voluntary isometric contraction reduced the magnitude of the evoked responses but not its spatiotemporal profile. Notably, TMS administered during the active motor state reliably attenuated the N15 and N100 TEP peak amplitudes across a broad range of stimulation intensities. This reduction was consistently observed across participants, underscoring the robustness of the effect.

The precise neural generators of the N15 and N100 TEP peaks and the mechanisms rendering these peaks sensitive to changes in motor state remain to be delineated [5]. Based on their latency, it can be excluded that the N15 and N100 peaks reflect the initial direct inductive activation of local cortical neurons responsible for generating the MEPs [30]. Instead, recent evidence suggests that immediate TEPs (iTEP), which emerge within 2-8 ms following the TMS pulse [13], may serve as a more direct indicator of immediate TMS-induced cortical activation [13]. In the present study, we were unable to assess iTEPs due to the presence of stimulation-related artifacts within the first 5-6 ms after the TMS pulse, due to the sampling rate of 5 kHz. Emerging evidence suggests that both early TEP peaks (e.g., N15) and late TEP peaks (e.g., N100) may instead reflect reverberant activity within cortico-thalamic-cortical circuits triggered by the TMS pulse [31–33]. Studies in mice and humans comparing electrical and magnetic stimulation during rest and movement have shown a consistent suppression of both early and late response amplitudes during movement [31,32]. The authors also performed a detailed mapping of the network responsible for mediating these effects. In mice, stimulation evoked early spiking in cortico-thalamic projection neurons, followed by sensorimotor thalamic activity, with this relationship reversing during later time windows [32]. Furthermore, optogenetic inhibition of sensorimotor thalamic neurons altered the cortical response profile [31]. Together, these findings support the notion that thalamic-cortical feedback is involved in shaping the TEP in a state-dependent fashion. While the precise timing and definition of the peaks in those studies differ from the ones reported in the present work, the cross-species characterization of movement-modulated circuits provides valuable insights into state-dependent modulation. We propose that the attenuation of N15 and N100 amplitudes by voluntary contraction may reflect the dynamic engagement of sensorimotor cortico-thalamic-cortical circuits. These TEP peaks may serve as sensitive markers of motor state transitions and potentially offer novel insights into motor dysfunction in neurological disorders. Further investigation is warranted to clarify their underlying mechanisms and clinical relevance.

### State-dependent effects of voluntary muscle activation on MEPs and TEPs

While MEP amplitudes were facilitated by voluntary contraction [34], the N15 and N100 peak amplitudes of the TEP were suppressed during tonic isometric contractions compared to rest. Preparation and execution of voluntary movement is characterized by a desynchronization (i.e., decrease of power) of cortical network activity at alpha (8-13 Hz) and beta (15-35 Hz) frequencies [23,35], and an increase in the excitability of spinal motoneurons. Cortical desynchronization likely reflects a transition from a synchronized oscillating state towards a state of more asynchronous neural processing that is needed to generate and maintain a tonic level of motor output. The greater MEPs following TMS of M1 during low intensity sustained voluntary contractions can be attributed to increased excitability of alpha motor neurons in the spinal cord [36]. Although generally representing a more active cortical state than rest, the evoked N15 and N100 TEP peaks were reduced during periods of isometric voluntary contraction relative to rest. This may be related to intracortical inhibition, primarily mediated by GABAergic interneurons, dampening excitatory post-synaptic activity. A potential contribution of cortical inhibitory circuits to the N15 and N100 peaks is supported by both pharmacological and paired pulse TMS studies [37–39]. Of note, a relative attenuation of the N100 peak has been found during both movement preparation [40–42] and execution [43,44]. The present results confirm and extend these N100 findings by showing that tonic isometric contraction also reduces the amplitudes of the early N15 peak. Together, it can be concluded that both early and late cortical responses to TMS are sensitive to the level of motor activity and attenuated by tonic voluntary motor activity. Since contraction-related attenuation of the N15 and N100 amplitude correlated in our participants, the mechanisms mediating state-dependent modulation of these two peaks may be at least partly overlapping. Alternatively, the attenuation could be generated by distinct mechanisms that are influenced to a comparable extent by tonic contraction.

It is worth noting that the interval between the N15 and N100 peaks roughly aligns with the alpha frequency band (8-13 Hz). In our study, voluntary tonic contraction of an intrinsic hand muscle led to a reduction in sensorimotor EEG power in both, the alpha and beta bands. However, individual decreases in alpha and beta power ipsilateral to stimulation did not correlate with the attenuation of N15 and N100 TEP components, nor with the individual facilitation of MEP amplitudes during tonic contraction. This lack of correlation suggests that individual changes in task-related regional cortical alpha and beta power do not account for the observed modulation of TEPs or MEPs. This finding is particularly relevant in light of recent work using combined EEG-TMS approaches to examine how oscillatory brain activity shapes TEP and MEP responses at rest. While several EEG-informed TMS-MEP studies have shown that MEP amplitudes are influenced by the power and phase of the pericentral alpha rhythm [45–48], similar effects have not been consistently demonstrated for TEPs [49,50].

### Dose dependency of MEP and TEP responses

MEPs were only elicited at threshold and suprathreshold intensities during rest, while MEPs were observed at all stimulation intensities during voluntary contraction, including the two intensities that were below the resting motor threshold. The latter suggest that the entire intensity range used in this study was above the threshold for evoking activity in pyramidal tract neurons. Accordingly, TMS over M1-HAND elicited EEG responses close to site of stimulation at all intensity levels, even in the subthreshold resting state conditions, evoking a well-described sequence of TEP peaks with latencies at 15, 30, 45, 60 and 100 ms [4,22,51]. While some peaks were clearly discernible, others were more difficult to disentangle such as the N45 and P60, as they were superimposed on a positive wave [52]. This effect seemed more pronounced following suprathreshold stimulation compared to subthreshold stimulation and likely also contribute to explaining the correlation of the state-dependent modulation between the two.

Increasing the intensity of TMS enhanced the response magnitude for both MEPs [34] and TEPs [10]. Stimulus intensities scaled positively with MEP and TEP amplitudes at rest and during a sustained voluntary contraction due to a more efficient excitation of cortical neurons. The excitation of a larger population of pyramidal cells at higher stimulus intensities should translate into larger amplitudes of both MEPs and TEPs. For TEPs, other non-transcranial stimulation effects become more pronounced at higher stimulation intensities. For example, the TMS-evoked muscle twitches in the contralateral hand or forearm muscles increase with stimulus intensity, causing stronger re-afferent somatosensory feedback that may increase middle-to-late parts of the TEPs (from approx. 40ms and onwards) [53,54]. Stronger stimulation intensities also produce greater mechanical vibrations in the coil casing which produce larger “click” sounds [55] that are more easily perceived by participants, as also reported in Table 1. Multimodal sensory inputs likely become more relevant at higher TMS intensities and contribute to the stimulus-response relationship of TEP components that occur at latencies above 40 ms through peripherally-induced cortical co-activation if not masked sufficiently [11,56–59].

**Table 1.** **Sensory perception of TMS pulses**.

|  | Loudness of pulse “click”# |  | Scalp “spread” of pulse |  | Coil pressure |  | Coil vibration |  |
| --- | --- | --- | --- | --- | --- | --- | --- | --- |
|  | Rest | Contraction | Rest | Contraction | Rest | Contraction | Rest | Contraction |
| -8% MSO | 1.46 ± 1.76 | 1.19 ± 1.72 | 5.87 ± 2.72 | 6.20 ± 2.65 | 1.87 ± 1.76 | 1.50 ± 1.26 | 0.13 ± 0.35 | 0.13 ± 0.34 |
| -4% MSO | 1.31 ± 2.09 | 1.50 ± 2.03 | 5.94 ± 3.02 | 6.31 ± 2.21 | 1.56 ± 1.36 | 1.69 ± 1.74 | 0.13 ± 0.34 | 0.31 ± 0.60 |
| RMT | 1.63 ± 1.93 | 1.69 ± 1.78 | 6.06 ± 2.67 | 6.31 ± 2.57 | 1.75 ± 1.53 | 1.63 ± 1.31 | 0.13 ± 0.34 | 0.25 ± 0.58 |
| +4% MSO | 1.86 ± 1.63 | 2.07 ± 1.65 | 6.06 ± 2.46 | 6.56 ± 2.25 | 1.56 ± 1.21 | 1.94 ± 1.39 | 0.31 ± 0.79 | 0.50 ± 1.21 |
| +8% MSO | 2.20 ± 2.57 | 2.80 ± 2.37 | 6.02 ± 2.48 | 6.40 ± 2.59 | 1.93 ± 1.79 | 1.87 ± 1.92 | 0.13 ± 0.35 | 0.33 ± 0.82 |

Sensations reported by the participants using a numerical rating scale (0-10). For loudness, 0 corresponded to not hearing the click sound, while 10 represented a very loud sensation. For focality, 0 represented a very focal sensation (i.e., “sharp”) and 10 very diffuse (i.e., “blunt”). For coil pressure, 0 represented no pressure at all and 10 very severe pressure. For coil vibration, 0 represented no vibration at all and 10 very intense vibration. Data reported as means and standard deviations. No significant differences between motor states were observed between subjective perceptions of auditory or somatosensory stimuli. # represents the significant effect of stimulation intensity across motor states for perceived loudness of the TMS coil “click”.

### Methodological considerations

Recording TEPs after stimulation of M1-HAND can be methodological challenging as the coil is placed on lateral aspects of the head in proximity to scalp muscles (e.g,, the temporal muscle) and the nerves that innervate them [1]. This increases the risk of coactivating scalp muscles resulting in large amplitude compound muscle action potentials that can obscure the EEG signal in the tens of milliseconds following stimulation [5,17]. Higher stimulation intensities are more likely to cause scalp muscle activation which further complicates mapping input-output relationships. By individualizing coil position through minor tilts and coil movements away from the temporal muscle [17] while concurrently visualizing the corresponding EEG data [16], we managed to acquire scalp muscle artifact free data in 16 out of screened 28 participants at a range of different intensities. In these individuals, we obtained short latency TEP responses (earliest peak at ∼15 ms) with amplitudes that resemble those seen after individualized and optimized targeting of peri-sagittal cortical areas (containing no muscles) [60].

We argue that our TEP responses were not confounded by scalp muscle activations. First, the amplitudes and topographical distribution do not resemble those of prototypical TMS-evoked scalp muscle artifacts [5,17,61]. Second, the fact that the N15 response was consistently modulated by the behavioral state of participants indicates that it is unlikely to represent scalp muscle artifacts which should not be altered by the central motor state manipulation.

In the present study, we compared TEPs during rest and sustained motor behavior. It is well established that sensory perception and attention are affected during motor activities [62–65]. For example, auditory evoked cortical potentials are smaller during periods of movement compared to rest [64,65]. In the present study, we used active noise masking to minimize the influence of auditory evoked potentials on the TEP [18,19]. Furthermore, no differences were observed in the subjective perception of any of the assessed sensory stimuli caused by TMS, including coil clicks. However, as we were not successful in completely masking the TMS-evoked click sound in all participants at all intensities, we cannot entirely exclude that the modulations of TEP peaks could reflect gating of sensory inputs caused by TMS (peripheral evoked potentials) or shifts in attention rather than a modulation of the transcranial constituent of the TEP per se. Peripherally evoked potentials primarily influence later TEP peaks [11,56–59]. Therefore, contributions of sensory gating of these potentials may be more relevant for the modulation of later (N100) rather than earlier (N15) TEP peaks.

## Conclusion

MEPs and TEPs are commonly used readouts of target engagement following stimulation of M1 and reflect the combined effect of the stimulation dose and the ongoing brain state. Here we show that the cortical responses (TEPs) and the cortico-motor pathway responses (MEPs) diverge when the physiological state switches from relaxation to tonic activation. The marked differences in state dependency show that TEPs and MEPs are distinct and do not offer two perspectives on the same physiological process. The state sensitivity of the classical TEP peaks (N15 and N100) may be a useful cortical probe of cerebral sensorimotor network dynamics in healthy individuals and patients.

## Acknowledgements

This study is funded by the Innovation Fund Denmark (IFD) for the Grand Solution project “PRECISION-BCT” (Grant number: 9068-00025B) and the project “ADAptive and Precise Targeting of cortex-basal ganglia circuits in Parkinson’s Disease - ADAPT-PD” from The Lundbeck Foundation (collaborative project grant, grant nr. R336-2020-1035). Mikkel M. Beck is funded by a grant from the Capital Region of Denmark (Region Hovedstaden) and a post doc grant from The Lundbeck Foundation (grant nr. R449-2023-1487). Sybren Van Hoornweder is funded by two grants from the Research Foundation Flanders (FWO) (Grant nr. G1129923N and nr. V426023N). Leo Tomasevic was partially funded by the dtec.bw – Digitalization and Technology Research Center of the Bundeswehr [MEXT project]. The dtec.bw is funded by the European Union – NextGenerationEU.

